# Identification of genetic markers for cortical areas using a Random Forest classification routine and the Allen Mouse Brain Atlas

**DOI:** 10.1101/549519

**Authors:** Natalie Weed, Trygve Bakken, Nile Graddis, Nathan Gouwens, Daniel Millman, Michael Hawrylycz, Jack Waters

**Affiliations:** Allen Institute for Brain Science, 615 Westlake Ave N, Seattle WA 98109

## Abstract

The mammalian neocortex is subdivided into a series of ‘cortical areas’ that are functionally and anatomically distinct, and are often distinguished in brain sections using histochemical stains and other markers of protein expression. We searched the Allen Mouse Brain Atlas, a database of gene expression, for novel markers of cortical areas. We employed a random forest algorithm to screen for genes that change expression at area borders. We found novel genetic markers for 19 of 39 areas and provide code that quickly and efficiently searches the Allen Mouse Brain Atlas.

## Introduction

The mammalian neocortex is classified into a series of anatomically and functionally distinct regions or ‘cortical areas’ (Brodmann, 1909; Glasser *et al.*, 2016). Areas are often identified using histochemical stains and antibodies to visualize differences in protein expression across cortex. Examples include cytochrome oxidase histochemistry and antibodies against m2 muscarinic receptors (Wang, Sporns & Burkhalter, 2012). Furthermore, global expression signatures of cortical areas have been identified in human (Hawrylycz et al., 2012), rhesus monkey (Bernard et al., 2012) and mouse (Hawrylycz et al., 2010), but few genes have been identified with distinct transitions between adjacent areas. We reasoned that there may be genetic markers of cortical areas that have not been identified and that we might identify additional markers by screening the Allen Mouse Brain Atlas, a database containing in situ hybridization information for thousands of genes (Lein *et al.*, 2007). We developed numerical tools to screen the many thousands of images in the database, using a random forest algorithm to identify changes in gene expression at the boundaries of cortical areas defined in the Allen Mouse Brain Reference Atlas (Kuan *et al.*, 2015). We found novel genetic markers for several areas. In addition, we provide code that searches the Allen Mouse Brain Atlas quickly and efficiently for differences in gene expression between cortical areas. With only minor modification, our code could be adapted to search for genes that mark other brain regions, including subcortical nuclei.

## Methods and Results

Our aim was to locate changes in gene expression between cortical regions in the mouse. From the Allen Mouse Brain Atlas, we took coronal in situ hybridization (ISH) data resampled to a canonical 3D reference space and overlaid the borders of cortical regions from the Allen Mouse Brain Reference Atlas. To identify genes with differential expression along these boundaries, we used a Random Forest algorithm.

### Horizontal Projections

We obtained ISH data for 4345 genes from the Allen Mouse Brain Atlas (brain-map.org/api/index.html). ISH data were of coronal sections (Figure 1A). However, the perspective that best captures most borders delineating cortical areas while eliminating excess information is the horizontal plane. To obtain a horizontal plane perspective from coronal sections, we created two projections for each gene: a ‘top projection’ and a ‘flat map projection’. Each projection was created in three steps, with the first two steps being common to both projections. Firstly, we isolated cortical fluorescence and eliminated fluorescence from subcortical structures by applying a mask derived from the Allen SDK (2015) structure_tree class (Figure 1B and C). Secondly, we created a maximum intensity surface projection: for each pixel on the cortical surface, we projected the fluorescence in the underlying tissue along a line perpendicular to the pial surface of cortex. One might think of this first step as creating a curved sheet of fluorescence intensity values at the surface of cortex. Finally, we projected these surface values to the horizontal plane, creating a top projection (Figure 1D) or we ‘unfurled’ the curved cortical sheet to create a flat map (Figure 1G). The flat map was particularly valuable in the study lateral cortical regions, which are under-represented in top projections.

**Figure 1.**
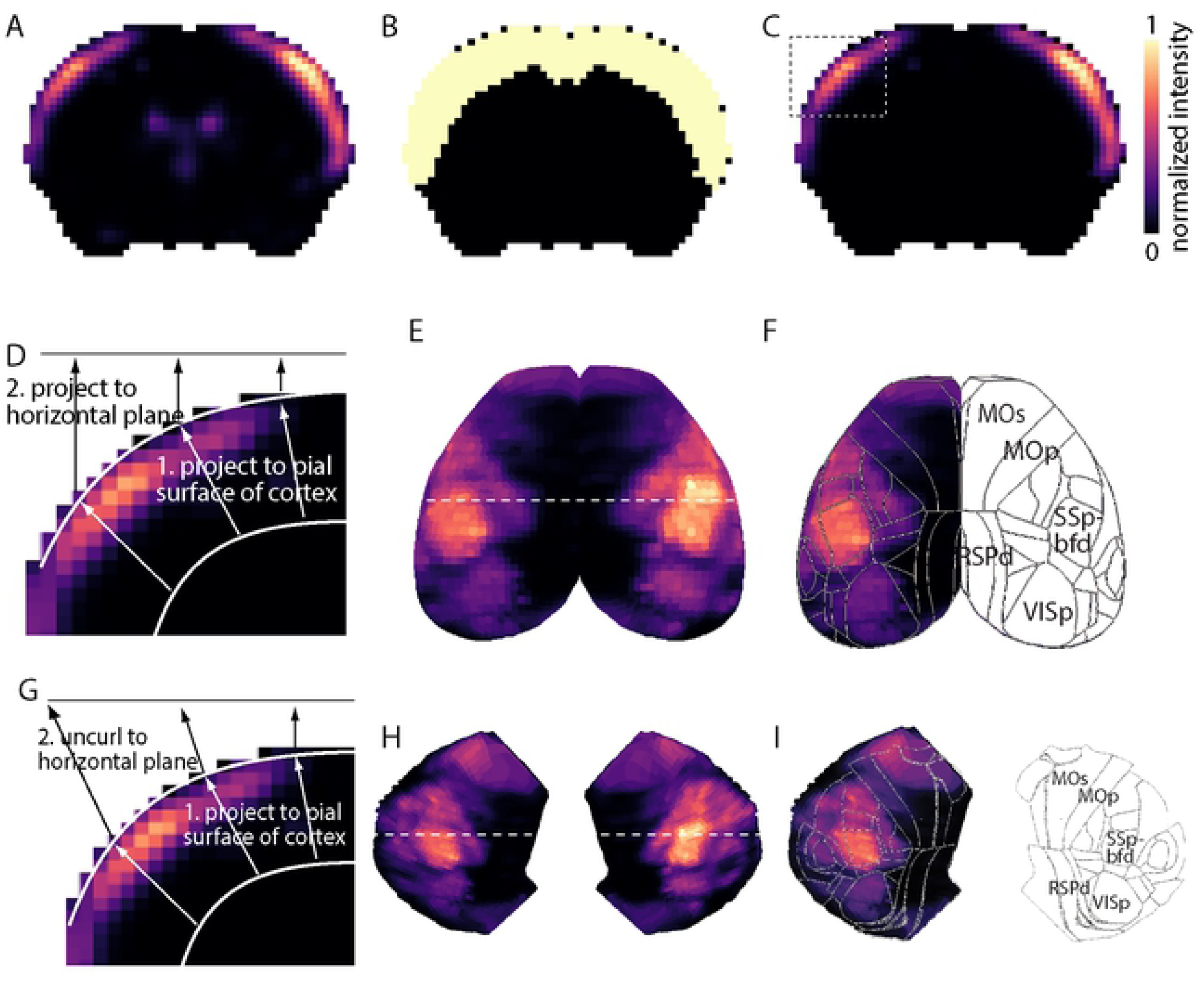
Creation of top projections from coronal images of gene expression. (A) Image of a coronal section from the Allen Mouse Brain Atlas. Gene: *Rorb*. ISH intensity is normalized to a range of 0 to 1. Color scale shown in panel C. (B) Binary cortex mask with value = 1 for cortical pixels and value = 0 for subcortical pixels. (C) Product of the images in A and B, resulting in ISH intensity values in cortical pixels and zero’s in subcortical locations. (D) Schematic illustration of the projection process used to generate top projections. (E) Top projection for *Rorb*. Dashed line indicates location of section in panel A. (F) Cortical boundaries from the Allen Mouse Brain Reference Atlas, overlaid onto the gene expression top view of panel E. (G) Schematic illustration of the projection process used to generate flat map top projections. (H) Flat map projection for *Rorb.* Dashed line indicates location of section in panel A. (I) Cortical boundaries from the Allen Mouse Brain Reference Atlas, overlaid onto the gene expression top view of panel H.

All ISH data in the Allen Mouse Brain Atlas are spatially registered to the Allen Mouse Brain Reference Atlas (http://help.brain-map.org/display/mousebrain/Documentation?preview=/2818169/8454277/MouseCCF.pdf). Hence all data utilized are inherently co-aligned with the Allen Mouse Brain Reference Atlas and the locations of brain areas can be readily superimposed on the ISH results. To locate cortical regions in the top projection and create a cortical area map, we extracted the corresponding cortical area masks using the structure_tree class and projected these masks to the horizontal plane, as described for ISH projections. Simplification of three-dimensional data into two dimensions allowed for fast quantitative analysis as well as easy visualization of expression patterns.

### Random Forest Algorithm

When examining the ISH results, two limitations became apparent. Firstly, there are gaps in some data sets, with missing data manifest as dark pixels in coronal images or dark medial-lateral bands in the top projections (Figure 2A). Secondly, there is pronounced section-to-section variability in mean fluorescence that appears as coronal banding or ‘stripes’ in top projections (Figure 2B). Together these two effects often result in variation in pixel values, independent of variation due to differential gene expression. These data properties complicate the comparison of fluorescence along the anterior-posterior axis and, thereby, the comparison of expression between cortical regions. Rather than attempt to mitigate these issues directly, we trained a Random Forest algorithm to classify pixels as either inside or outside each cortical region, essentially learning the variance in the data.

**Figure 2.**
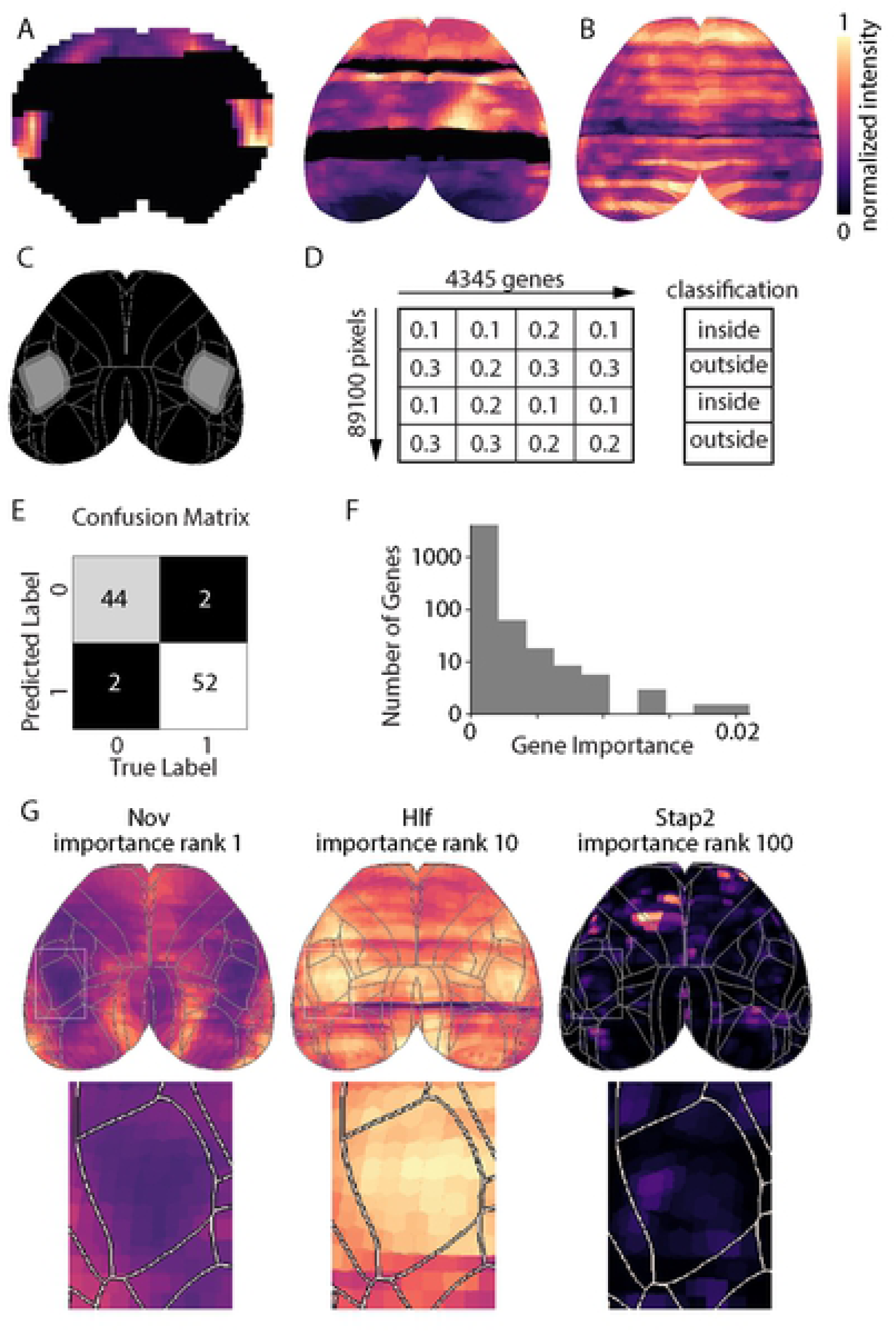
Random forest algorithm: method and example results. (**A**) A coronal image (left) and the top projection (right) for gene *Nvl*. Note the missing data (black pixels). (**B**) Top view for gene *Adra1d*. Note the pronounced variation in density along the a-p axis. (**C**) Binary mask for primary somatosensory cortex barrel field (SSp-bfd). Light gray inside SSp-bfd; darker grey marks pixels in surrounding region. White lines: boundaries of cortical areas in Allen Mouse Brain Reference Atlas. (**D**) Schematic illustration of arrays input into Random Forest algorithm. Columns correspond to gene, rows to pixels in the top projection data set. Each value is an ISH luminance value. Classification of pixel is taken from the reference mask (panel C). (**E**) Confusion matrix output from Random Forest algorithm for SSp-bfd. 0 indicates point outside SSp-bfd, 1 indicates point inside SSp-bfd. (**F**) Gene importance histogram. Importance values approximate a logarithmic distribution. (**G**) Examples of genes that mark SSp-bfd, with overlaid Allen Mouse Brain Reference Atlas borders. *Nov* rank 1, importance 0.022. *Hlf* rank 10, importance value 0.0081. *Stap2* rank 100, importance 0.0018.

We examined 39 cortical regions from the Allen Mouse Brain Reference Atlas for potential gene markers. Each search involved comparison of one cortical region to expression patterns of all genes, imputed as independent variables to the Random Forest algorithm. Random forest was implemented in Python using the scikit-learn package (Pedregosa *et al.*, 2011). Nodes were determined by Gini Index criteria 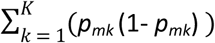. Each random forest consisted of 100 decision trees. Random state was initialized at 0. The importance of each variable was also determined by Gini Index criteria – reduction in Gini Index each time a split occurred was attributed to the variable, and that variable-associated reduction was divided by total reduction in Gini Index across the entire random forest to return the variable importance value. Total variable importance across all genes summed to one.

For each cortical region, three outputs from the random forest were produced and analyzed: (1) a confusion matrix, indicating the success rate of the classification algorithm; (2) the list of all 4345 genes, ranked in decreasing order of variable importance, where importance is a pseudo-measure of the expression predictive power across the cortical border; and (3) the importance values, one for each gene. The Random Forest was trained to distinguish pixels within a cortical region from pixels in the surrounding area outside of the cortical region. The inputs to the Random Forest algorithm were the gene expression fluorescence intensity values of all 4345 genes for each pixel and the corresponding labels for each pixel as cortical region or surrounding region. Surrounding pixels were identified by dilating the region mask by 30 iterations using SciPy ndimage package in Python, translating to roughly 30 pixels in distance in each direction. Pixels were split into training and test sets, with 100 randomly selected pixels held out as a test set and the remaining pixels forming the training set. This represented less than 1% of total pixels classified for each cortical area. Data was divided using the scikit-learn model_selection package in Python. Hence the training array input into the Random Forest algorithm consisted of a 2D array of dimensions 4345 by *N* ‒ 100, where 4345 is the number of genes and *N* is the number of pixels within the dilated mask, and each cell in the array corresponding to a luminance value of the pixel. A second array of dimensions 1 by *N* ‒ 100 indicated the binary labels, inside or outside the cortical region (Figure 2D). After training, performance of the algorithm was tested on the held-out pixels (array dimensions 4345 by 100) for which the binary classification was withheld. Withheld pixels were randomly selected, creating a test set that was representative of the cortical area: balanced inside and outside the cortical region, and varying in distance from the cortical area boundary. Hence the algorithm returned the cross-validated binary classification for 100 withheld pixels, which was compared to known classification and used to plot a confusion matrix (Figure 2E), summarizing performance of the Random Forest. The displayed confusion matrix is averaged over all folds for the specified cortical region.

Results for primary somatosensory barrel field are illustrated in Figure 2E-G. The model correctly classified 52 of 54 test pixels within the barrel field and 44 of 46 test pixels outside barrel field, resulting in a combined model accuracy of 96% (Figure 2E). Most genes exhibited low variable importance (Figure 2F). We ranked genes by their random forest variable importance values. The gene with rank 1 exhibited a distinct change along the border (Figure 2G). The gene with rank 10 exhibited a subtler change and the gene at rank 100 exhibited no obvious change along the border (Figure 2G). Hence the Random Forest algorithm accurately classified most pixels and, via a ranked list of genes, identified a short list of genes that might act as putative genetic markers of the cortical region.

### Genetic markers of cortical areas

To identify genetic markers for each cortical area, we manually inspected the top projections or flat maps for the top 10 genes, as determined by the Random Forest results. Adequate information for classification of barrel field was included in the 10 highest importance genes since running our analysis with only these 10 genes as inputs conserved prediction accuracy at 96%. Of the 45 cortical regions tested, we identified potential genetic markers for 19 (Table 1). 6 cortical areas were determined too small to reliably examine, resulting in 39 regions to explore. Of the six markers identified by Hawrylycz *et al.* (2010), three were extracted by our method (*Man1a*, somatomotor; *Rorb*, somatosensory; *Scnn1a*, ventral retrosplenial), one included an area that was not explored (*Smoc1*, gustatory), one was in a region where many other strong markers were identified (*Rreb1*, retrosplenial), and one was not identified (*Hap1*, ectorhinal). Selected potential genetic markers indicated relatively high sensitivity for area marking, low specificity due to the point selection process, bilateral expression, and entire cortical area contrast.

**Table 1.**
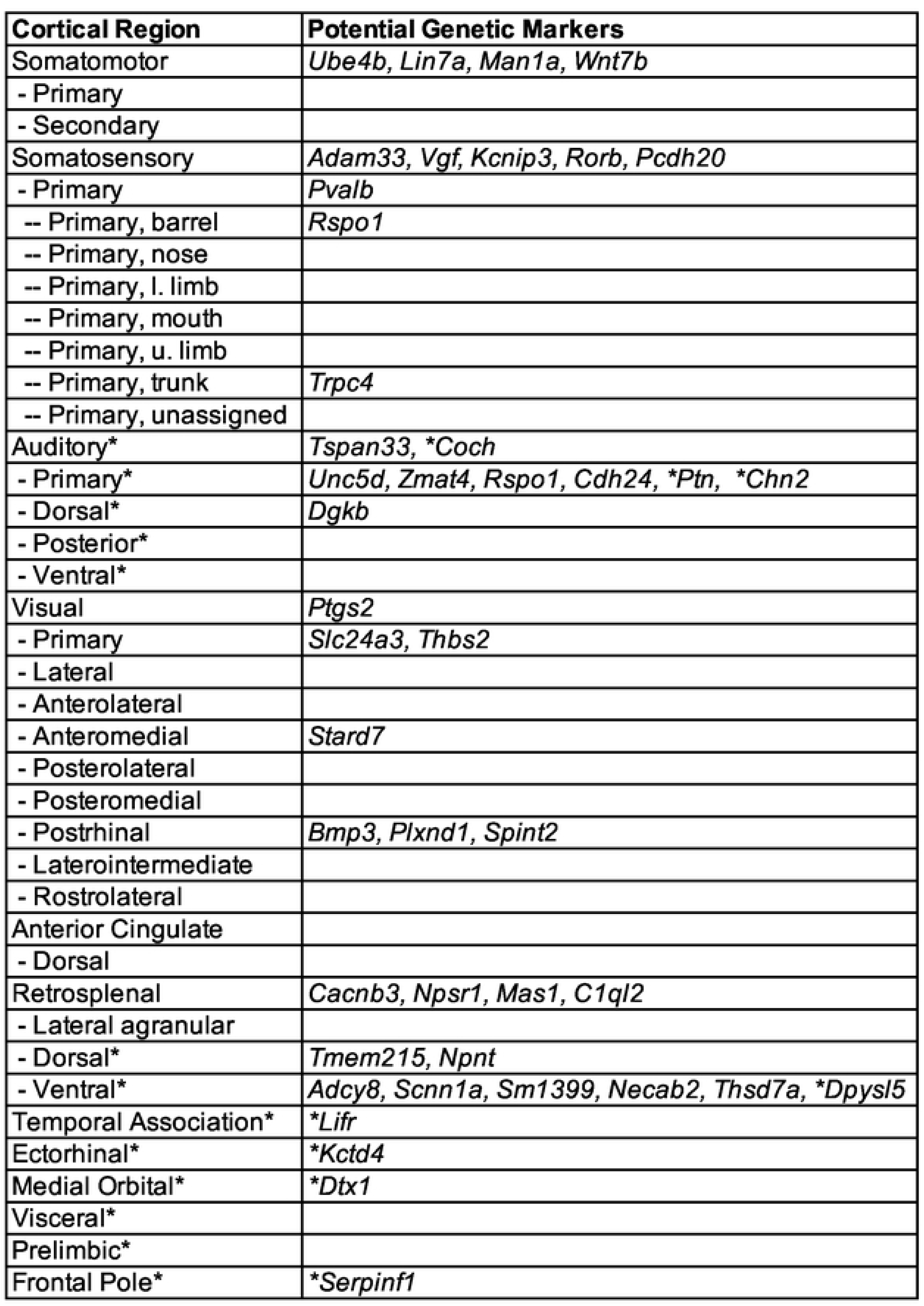
List of Potential Genetic Markers. Potential cortical boundary genetic markers, listed by cortical region, as identified by Random Forest variable importance classifier. All explored regions listed. Regions with no listed genes displayed no clear potential genetic marker. Each gene was identified independently. Asterisks indicate regions explored and genes identified with flat map projections.

| **Cortical Region** | **Potential Genetic Markers** |
| --- | --- |
| Somatomotor | *Ube4b, Lin7a, Man1a, Wnt7b* |
| - Primary |  |
| - Secondary |  |
| Somatosensory | *Adam33, Vgf, Kcnip3, Rorb, Pcdh20* |
| - Primary | *Pvalb* |
| -- Primary, barrel | *Rspo1* |
| -- Primary, nose |  |
| -- Primary, l. limb |  |
| -- Primary, mouth |  |
| -- Primary, u. limb |  |
| -- Primary, trunk | *Trpc4* |
| -- Primary, unassigned |  |
| Auditory* | *Tspan33, \*Coch* |
| - Primary* | *Unc5d, Zmat4, Rspo1, Cdh24, \*Ptn, \*Chn2* |
| - Dorsal* | *Dgkb* |
| - Posterior* |  |
| - Ventral* |  |
| Visual | *Ptgs2* |
| - Primary | *Slc24a3, Thbs2* |
| - Lateral |  |
| - Anterolateral |  |
| - Anteromedial | *Stard7* |
| - Posterolateral |  |
| - Posteromedial |  |
| - Postrhinal | *Bmp3, Plxnd1, Spint2* |
| - Laterointermediate |  |
| - Rostrolateral |  |
| Anterior Cingulate |  |
| - Dorsal |  |
| Retrosplenial | *Cacnb3, Npsr1, Mas1, C1ql2* |
| - Lateral agranular |  |
| - Dorsal* | *Tmem215, Npnt* |
| - Ventral* | *Adcy8, Scnn1a, Sm1399, Necab2, Thsd7a, \*Dpysl5* |
| Temporal Association* | *\*Lifr* |
| Ectorhinal* | *\*Kctd4* |
| Medial Orbital* | *\*Dtx1* |
| Visceral* |  |
| Prelimbic* |  |
| Frontal Pole* | *\*Serpinf1* |

Examples of expression patterns are provided in Figure 3. For primary somatosensory cortex barrel field, we identified *Rspo1* as a strong candidate gene (Figure 3A). Expression of *Rspo1* is relatively high in the barrel field, moderate through somatosensory areas, and low in motor cortex. There were multiple markers for motor cortex, including *Wnt7b* (Figure 3B), but we found no compelling markers for primary or secondary motor cortex. *Rorb* was also identified as a potential marker, specifically for primary sensory cortices (Figure 3C). This provided an additional positive control that our method was robust and effective, as *Rorb* is an established marker for primary sensory areas (Hawrylycz *et al.*, 2010; Zhuang *et al.*, 2017). *Cdh24* marked primary auditory cortex (Figure 3E). We found multiple genes that labeled all or subregions of retrosplenial cortex. For example, *Tmem215* marked dorsal retrosplenial cortex (and primary somatosensory cortex) and *Npsr1* marked all of retrosplenial cortex (Figure 3D, F). In flat maps, *Serpinf1* was identified as a marker of the frontal pole (Figure 3G) and temporal association cortex was marked by *Lifr* (Figure 3H). Notably, some of these genes exhibit mediolateral stripes in the top projection, indicating that our method is robust to the missing data and expression-independent variability in signal.

**Figure 3.**
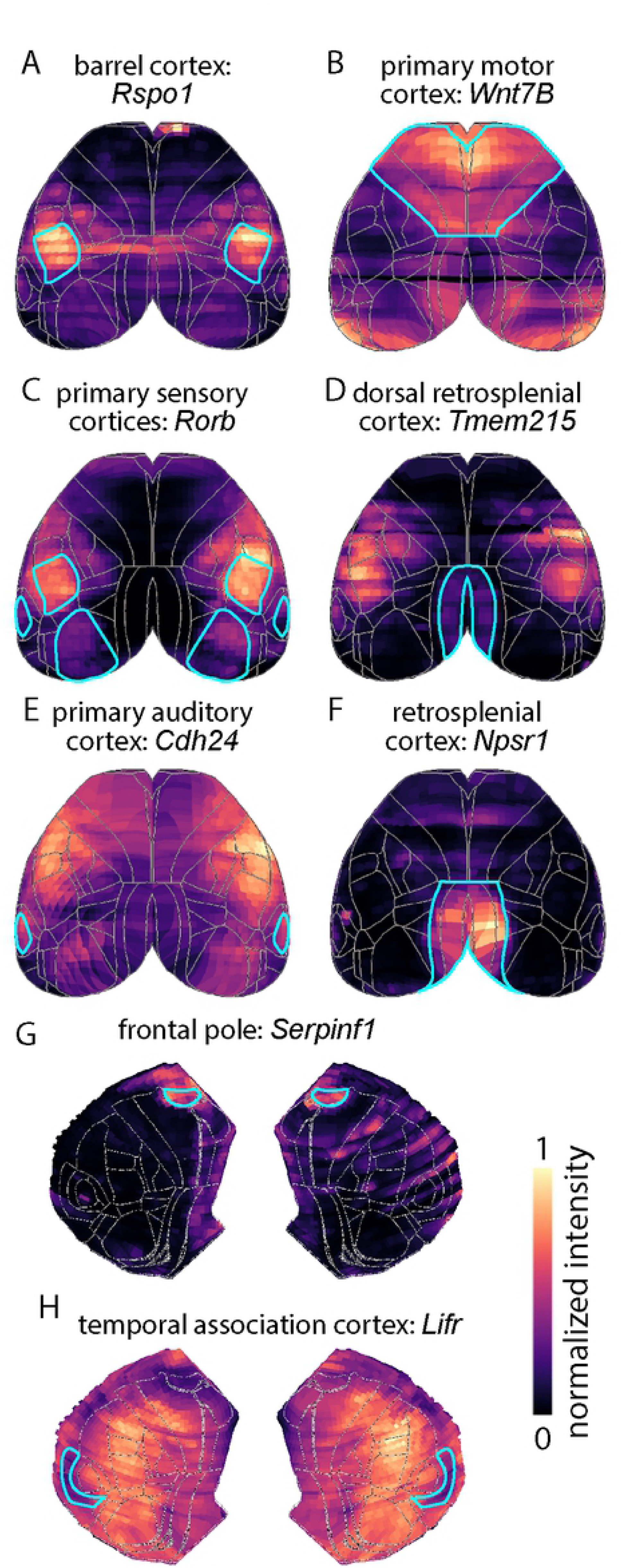
Examples of markers for cortical areas. R-Spondin 1 labels primary somatosensory cortex barrel field. (B) Wnt Family Member 7B labels primary motor cortex. (C) Retinoid-Related Orphan Receptor, Beta labels primary sensory areas. (D) Transmembrane Protein 215 labels dorsal retrosplenial cortex. (E) Cadherin 24 labels primary auditory cortex. (F) Neuropeptide S Receptor 1 labels retrosplenial cortex. (G) Serpin Family F Member 1 labels frontal pole. (H) Leukemia Inhibitory Factor Receptor labels temporal association cortex. Panels A-F provide examples of genes identified in top projections, panels G and H of genes identified in flat maps. Cortical areas of interest are marked with cyan borders.

### Cellular Basis of Potential Genetic Markers

Our Random Forest searches, applied to ISH data, identified genes that marked cortical area borders, but provided no insight into the cellular basis of the genetic markers. Does, for example, a change in gene expression result from an abrupt change in the density of a cell type with unique gene expression; or might the border result from a change in gene expression by a cell type that straddles the border? Gene expression in primary visual cortex and anterior lateral motor cortex has been studied using single-cell RNA sequencing (Tasic *et al*, 2018). From this transcriptomic data set and associated analysis tools (https://github.com/AllenInstitute/scrattch.vis), we examined cell types that express the marker genes identified in this study by top view projections (Figure 4). Markers were expressed in many different cell types. Several genes (*Adcy8, Bmp3, Cacnb3, Npsr1, Vgf, Zmat4*) were expressed mostly in neurons and not non-neuronal cells, suggesting that the border-related change in expression of these genes was neuronal. Some genes were expressed mostly in a cell sub-population, suggesting that there is likely a border-related change in the density of these cells or of their expression of one gene. For example, *Rspo1*, *Serpinf1* and *Man1a* are expressed in layer 4 excitatory neurons, vascular and leptomeningeal cells (VLMC) and macrophages, respectively. Unsurprisingly, our results are consistent with changes in gene expression marking cortical areas arising from changes in cell density in some instances and from changes in gene expression within a cell population in other instances. Importantly, both instances appear to have been detected by our Random Forest analysis of ISH data.

**Figure 4.**

Single Cell RNA-sequencing expression plot. Log-transformed average expression of top-projection identified potential areal marker genes in mouse cortical cells grouped into subtypes of three major cell classes: inhibitory neurons, excitatory neurons, and non-neuronal cells. Expression data was measured by RNA-sequencing of single cells isolated from Primary Visual Cortex (VISp) and Anterior Lateral Motor Cortex (ALM). Color corresponds to expression value, with warmer colors indicating high expression and cooler colors indicating low expression. Max value indicates the maximum expression per gene, measured in average CPM per cluster. CPM = counts per million reads.

## Discussion

We used a Random Forest algorithm to identify a short list of potential gene markers from thousands of candidate genes, applying this approach to 39 cortical regions in the mouse. Our results identified 44 putative markers, marking 19 of the explored regions.

The spatial resolution and number of genes in the database places limits on the conclusions we can draw. Firstly, the voxel size of the ISH quantification in the database is 200 µm. Once missing and variable data is considered, the maximum accuracy we can hope to achieve is on the order of hundreds of micrometers, resulting in an imperfect match between area borders and gene expression. Subsequent experiments such as immunostaining for the genes we have identified would be necessary to confirm our results and to assess the accuracy with which each gene marks borders. Furthermore, the database includes coronal ISH images for 4345 genes. It may be that genes not sampled here mark some of the 25 cortical regions for which we were unable to identify markers. Repeating our analysis on a larger data set, should one become available, might identify further markers.

Alternative methods include gene identification by direct comparison of expression difference along the cortical border. However, pooling of more pixels than those available solely along borders was necessary to overcome high luminance variability across pixels and coronal sections. For this reason, and the difficulty of direct quantification of variance in our data set, we decided to pursue random forest classification as our selected model. Variable importance may be inaccurately skewed towards higher sampled variables or continuous data types, and thus unusable; however, because our predictor variables exhibit identical scale of measurement and data type, importance rank can be taken as unbiased (Strobl *et al.*, 2007). Random forest uses a bootstrapped subset of variables at each splitting node when building decision trees. By accumulating many splits on previously subdivided pixels, genes are evaluated at subregions of the cortical area. Given this property, we find that occasionally genes with relatively high variable importance display marking of a single border rather than the entire cortical area. However, if a gene exhibits clear marking of all cortical borders, it is shown with higher variable importance than an alternate gene expression pattern marking only a single border. Random forest is an accurate, computationally efficient, and easily interpretable method of classification. This was important as many of our data sets, especially for larger cortical areas like somatosensory areas, reached sizes of almost 250,000 pixels, evaluated at 4345 genes. Each output predicted took less than a minute to compile the data set, run computations, and produce outputs on a desktop computer. By maximizing concurrent computation across all available cores, the time required to run is minimized while not sacrificing predictive power of our model, as exhibited by the high accuracy of the random forest.

By dilating the cortical area mask a small amount instead of comparing the area of interest to the entire cortex, we allowed for differential expression of the gene in more distant parts of cortex. This is by design, as expression far from the desired cortical region does not impact the ability of the gene to mark the border. However, potential uniquely expressed genes are still a subset of those that can be identified with our method, and our method could be readily modified to solely identify uniquely expressed genes. Similarly, the method could be easily extended to investigate laminar differences or expression patterns in subcortical structures as masks for cortical layers and for subcortical structures are included in the Allen Mouse Brain Reference Atlas.

## Acknowledgements

We thank the Allen Institute founder, Paul G. Allen, for his vision, encouragement and support.

